# Quantification of the necessary labelling input in protein stable isotope probing

**DOI:** 10.1101/2020.06.02.129254

**Authors:** Robert Starke

## Abstract

Even though protein stable isotope probing is at least 100-times more sensitive than nucleic acid approaches, it is still unknown how much labelling input is necessary to not lose incorporation. Here, I used the average peptide model together with primary incorporation rates from previous experiments to determine the minimally necessary labelling input to detect incorporation of atoms present in proteins. Carbon-13 showed the highest efficiency for incorporation with detection misses only below 20.24 atom% labelling input. Despite the highest abundance in proteins, deuterium incorporation is not lost with at least 32.22 atom% labelling input due to instable acidic protons that undergo HD-exchange. All incorporation of oxygen-18 is detected with at least 28.15 atom% labelling input due to the +2 mass shift between its isotopes making up for its low abundance in proteins. Nitrogen-15 is less abundant in proteins and therefore harbors the most inefficient incorporation, resulting in a minimally necessary labelling input of 73.10 atom% labelling input. Altogether, these results can be used to properly design stable isotope probing experiments with the aim to maximize incorporation that can be extrapolated to other nucleic acid approaches due to similar incorporation routes.

---

Metaproteomics, a central tool of microbial ecology (Bastida *et al*., 2009; Von Bergen *et al*., 2013), provides the direct link between phylogeny and function in microbial communities (Chourey *et al*., 2010; Keiblinger *et al*., 2012; Bastida *et al*., 2015; Starke, Jehmlich and Bastida, 2018) but metabolic activity of an organism within the community can only be unveiled by the application of protein stable isotope probing (SIP) (Jehmlich *et al*., 2008, 2010; Bastida *et al*., 2010). Stable isotopes of carbon (as carbon-13), hydrogen (as deuterium), nitrogen (as nitrogen-15), oxygen (as oxygen-17 or oxygen-18) and sulfur (as sulfur-33, sulfur-34 or sulfur-36) could potentially be traced into proteins. Of those, carbon-13 is commonly used for protein-SIP (Jehmlich *et al*., 2008; Taubert *et al*., 2012; F. A. Herbst *et al*., 2013; Starke, Keller, *et al*., 2016) with the aim to trace the degradation of single compounds and of complex mixtures. The decomposition of nitrogen-15-labelled tobacco was followed in proteins of a soil microbial community (Starke, Kermer, *et al*., 2016), sulfur-34 into amino acids (F.-A. Herbst *et al*., 2013) as well as deuterium (Taubert *et al*., 2018; Starke *et al*., 2020) and oxygen-18 (Starke *et al*., 2020) from heavy water as activity tracer in groundwater microbes. Otherwise, oxygen-17, sulfur-33 and sulfur-36 have not yet been utilized in protein-SIP, presumably due to the high cost of labelled material but also the low abundance of sulfur in biomolecules. However, stable isotopes possess the ability to highlight metabolic activity of an organism within a complex microbial community (Jehmlich *et al*., 2008, 2010; Bastida *et al*., 2010), especially in environments with a high degree of dormancy. Logically, the most efficient application is of greatest interest to govern future research but little is known about the necessary labelling input, which I aim to calculate here. I hypothesize different minimally necessary labelling inputs for the different atoms present in a protein due to differences in their abundance in proteins and their mass between the light and heavy isotope. I focused on carbon-13, deuterium, nitrogen-15 and oxygen-18 because isotopes of sulfur are practically impossible to use for protein-SIP (F.-A. Herbst *et al*., 2013) and oxygen-17 is extremely rare (Lide, 2005) and thus, very expensive.

Before the analysis of incorporation, one has to understand the outcome of protein-SIP. Typically, a Gaussian distribution of incorporation is detected in the spectrum (**Figure 1**). The monoisotopic peak depicted at 0.00 (or 0%) is followed by several and most commonly three peaks each with a mass difference of one (depicted as +1, +2 and +3), originating from the natural abundance of heavy stable isotopes (Lide, 2005). Logically, the incorporation pattern of the experiment must have maxima of the Gaussian distribution beyond the natural abundance pattern to be detected. For this reason, I further considered the +4 mass as minimally possible maxima of the isotopic pattern that can be detected, which accounts for four (in case of carbon-13, deuterium and nitrogen-15) or two (in case of oxygen-18) incorporated isotopes depending on the mass difference between the light and the heavy isotope. In addition, the lower detection limit was previously determined to be 0.01 for carbon-13 (Von Bergen *et al*., 2013) and is likely in a similar range for the other isotopes present in proteins. Otherwise, the upper detection limit of the relative isotope abundance (RIA) is typically around 0.9 since higher values almost always originate from the natural abundance pattern of a new peptides and hence, a false positive detection of incorporation. In SIP, theoretically, if 100 atom% input material is entirely consumed and incorporated into the biomass of the organism(s), one would expect complete incorporation with only one peak at a RIA of 1.0 (or 100%). Complete incorporation is generally not achieved since (i) organisms typically have different sources of every atom in the protein, i.e. when degrading an entirely carbon-13-labelled hydrocarbon, carbon-13 from CO_2_ could be incorporated to dilute the atom% of the input material (Taubert *et al*., 2012; Starke, Keller, *et al*., 2016), (ii) cells in natural systems are in different states of growth and will therefore incorporate isotopes at a differential efficiency and (iii) the incubation times are too short to achieve complete incorporation, or any combination of these. The incorporation rates as maximum of the Gaussian distribution compared to the labelling input as RIA value were previously determined to be 0.5 (or 50%) for carbon-13 (Taubert *et al*., 2012), 0.2 (or 20%) for deuterium (Taubert *et al*., 2018; Starke *et al*., 2020), 0.6 (or 60%) for nitrogen-15 (Starke, Kermer, *et al*., 2016) and 0.6 (or 60%) for oxygen-18 (Starke *et al*., 2020). The relative difference between the monoisotopic peak at 0.00 (or 0%) and the maximum of the Gaussian distribution is explained as labelling ratio (LR). A successful protein-SIP application yields LR values of more than 0.00 (or 0%), which would mean no incorporation from the labelling experiment referred to as underlabelling, and below 1.00 (or 100%), which would mean no natural abundance pattern referred to as overlabelling. However, the natural abundance pattern is necessary for both the identification of the peptide and the quantification of the isotope incorporation (Sachsenberg *et al*., 2015). Therefore, prior knowledge on the time of the process of interest is important to avoid both under- and overlabelling. Both the addition of unlabelled controls and the *de novo* identification of peptides without monoisotopic peaks represent two so far unsuccessful ways to reduce the loss of incorporation. In general, to circumvent overlabelling, I advise for short incubation times since isotope incorporation into proteins appears to be fast and without changes in RIA with prolonged incubation (Jehmlich *et al*., 2008; Taubert *et al*., 2012, 2018; Starke, Jehmlich and Bastida, 2018). Differently, a pulse of additions could be used instead of an one-time application of input material (Starke, Keller, *et al*., 2016) that increase carbon use efficiency and therefore not only avoid overlabelling but also excessive respiration through lower priming.

**Figure 1:**
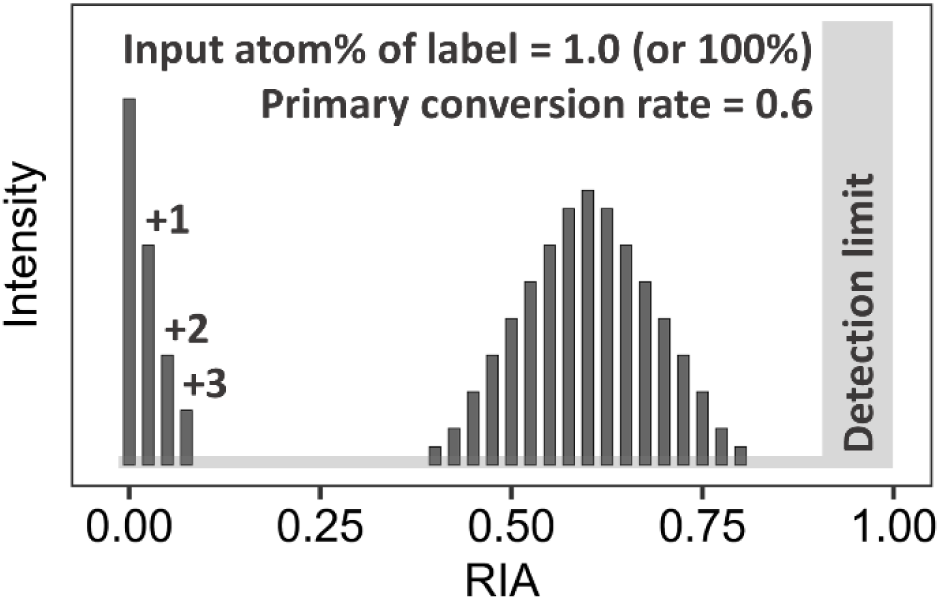
Gaussian distribution of a representative isotope incorporation into a peptide as relative isotope abundance (RIA) with a mass difference of +1 and a primary conversion rate of 0.6 of the input material, resulting in a maximum of 0.6 of the incorporation pattern. The lower detection limit depends on the mass difference between heavy and light isotope in each peptide. Otherwise, the upper detection limit is roughly 0.9 for all atoms.

Consistent with my hypothesis, carbon-13 showed the best efficiency of detecting incorporation when at least 20.24 atom% input label are supplied, resulting from the high number of carbon atoms in proteins (**Figure 2**). With a higher abundance in proteins, deuterium showed less incorporation efficiency requiring at least 32.22 atom% of input material because of the low incorporation rate of only 20% (Taubert *et al*., 2018; Starke *et al*., 2020). This is because of the loss of acidic deuteria in the HD-exchange (Molday, Englander and Kallen, 1972; Bai *et al*., 1993) once in contact with water during sample preparation and measurement. Even though both sample preparation and measurement could be performed in similar atom% of label, the amount would have to be adjusted to the number of acidic protons since every C-D bond is stable; a nearly impossible task especially in complex mixtures of proteins. In a similar range than deuterium, oxygen-18 required at least 28.15 atom% of input material. Despite its low abundance in proteins, the mass difference of two between the light and the heavy isotope allows for the detection of an average incorporation of only two isotopes and thus, a lower minimally necessary labelling input. Otherwise, the low abundance of nitrogen-15 in proteins together with a mass difference of one between the heavy and light isotope resulted in at least 73.10 atom% of input material labelling in order to avoid losing incorporation. Altogether, these results quantify the necessary labelling input in protein-SIP, which can easily be transferred to nucleic acid SIP approaches due to similar incorporation pathways into the biomass of the organisms when the minimally detectable incorporation is considered.

**Figure 2:**
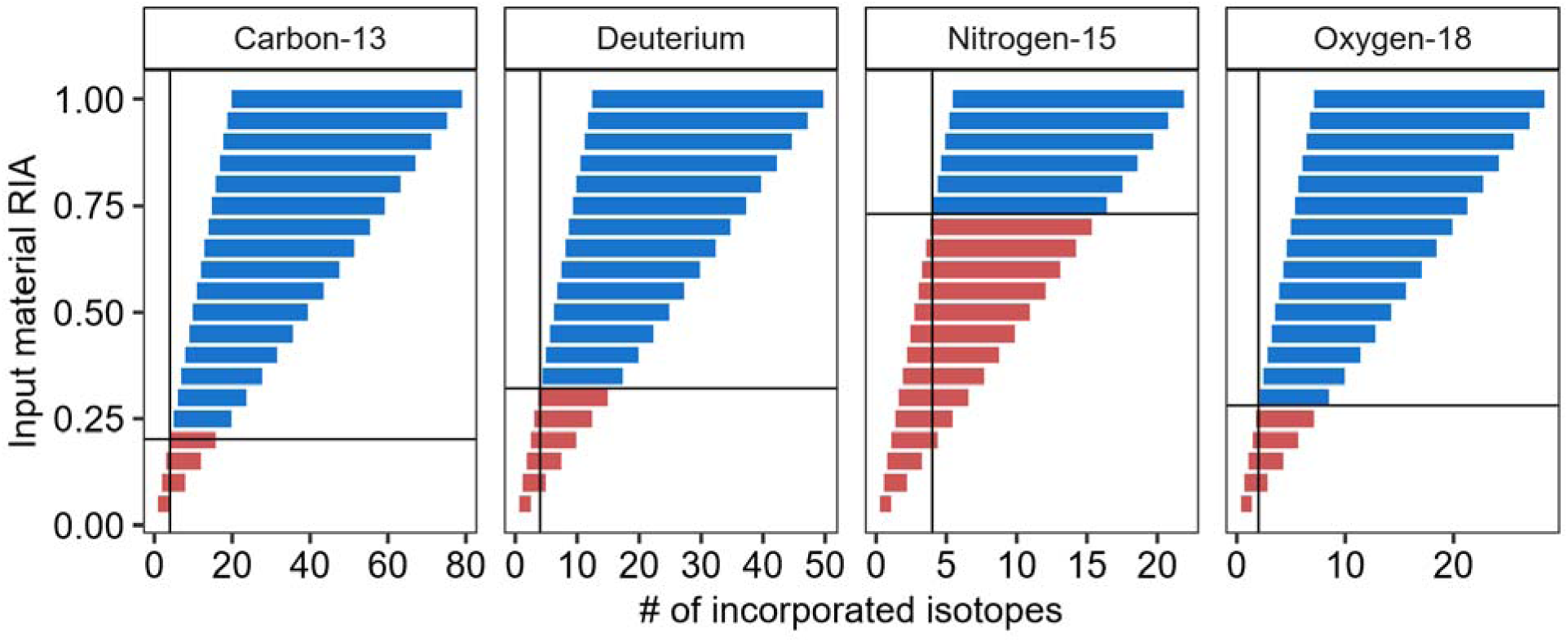
The number of isotopes incorporated into the typical peptide range from 8 to 32 amino acids relative to the input material relative isotope abundance (RIA) in intervals of 0.05 (or 5%) using the primary conversion rate of isotopes (0.5 for carbon-13, 0.2 for deuterium, 0.6 for nitrogen-15 and 0.6 for oxygen-18). The lower detection limit is indicated by the vertical line (4 for carbon-13, deuterium and nitrogen-15, and 2 for oxygen-18) due to the mass difference of 2 between oxygen-16 and oxygen-18. The minimally necessary input material RIA to avoid the loss of incorporation is depicted as horizontal line. Red boxes indicate input material RIA where a loss of incorporation can be expected.

## Materials and Methods

Using the average model of an amino acid of C_4.94_H_7.76_N_1.14_O_1.48_S_0.04_ (Senko, Beu and McLaffertycor, 1995), the number of incorporated isotopes for C, H, N and O of expected peptides (*n*_*isotopes*_) from the mass spectrometric measurements ranging from 8 to 32 amino acids (*n*_*aa*_) was estimated in steps of 5% with respect to the atom% of the input material (*RIA*_*input*_) and the primary incorporation rate *a*_*1st*_ (carbon-13 = 0.5 (Taubert *et al*., 2012), deuterium = 0.2 (Taubert *et al*., 2018; Starke *et al*., 2020), nitrogen-15 = 0.6 (Starke, Kermer, *et al*., 2016) and oxygen-18 = 0.6 (Starke *et al*., 2020)) according to Equation 1.

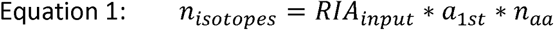

Given the representative incorporation pattern expected from protein-SIP that shows naturally abundant isotopes at the positions +1, +2 and +3 to the monoisotopic peak (**Figure 1**), I estimated the minimally detected incorporation (RIA as maximum of the Gaussian distribution) to be four incorporated isotopes (or +4 as mass) in case of carbon-13, deuterium and nitrogen-15. For oxygen-18, otherwise, the minimally measured incorporation is two incorporated isotopes (or +4 as mass) due to the mass difference between the oxygen-16 and oxygen-18. Combined, the expected RIA and the minimally possible detected RIA can be used to conclude necessary labelling input to yield efficient incorporation (**Figure 2**).

## References

Bai, Y. et al. (1993) ‘Primary structure effects on peptide group hydrogen exchange’, Proteins: Structure, Function, and Bioinformatics. doi: 10.1002/prot.340170110.

Bastida, F. et al. (2009) ‘Soil metaproteomics: A review of an emerging environmental science. Significance, methodology and perspectives’, European Journal of Soil Science. doi: 10.1111/j.1365-2389.2009.01184.x.

Bastida, F. et al. (2010) ‘Elucidating MTBE degradation in a mixed consortium using a multidisciplinary approach’, FEMS Microbiology Ecology. doi: 10.1111/j.1574-6941.2010.00889.x.

Bastida, F. et al. (2015) ‘Deforestation fosters bacterial diversity and the cyanobacterial community responsible for carbon fixation processes under semiarid climate: A metaproteomics study’, Applied Soil Ecology. doi: 10.1016/j.apsoil.2015.04.006.

Von Bergen, M. et al. (2013) ‘Insights from quantitative metaproteomics and protein-stable isotope probing into microbial ecology’, ISME Journal. doi: 10.1038/ismej.2013.78.

Chourey, K. et al. (2010) ‘Direct cellular lysis/protein extraction protocol for soil metaproteomics’, Journal of Proteome Research. doi: 10.1021/pr100787q.

Herbst, F.-A. et al. (2013) ‘Sulfur-34S Stable Isotope Labelling of Amino Acids for Quantification (SULAQ34) of Proteomic Changes in Pseudomonas fluorescens during Naphthalene Degradation’, Molecular & Cellular Proteomics. doi: 10.1074/mcp.m112.025627.

Herbst, F. A. et al. (2013) ‘Elucidation of in situ polycyclic aromatic hydrocarbon degradation by functional metaproteomics (protein-SIP)’, Proteomics. doi: 10.1002/pmic.201200569.

Jehmlich, N. et al. (2008) ‘Protein-based stable isotope probing (Protein-SIP) reveals active species within anoxic mixed cultures’, ISME Journal. doi: 10.1038/ismej.2008.64.

Jehmlich, N. et al. (2010) ‘Protein-based stable isotope probing’, Nature Protocols. doi: 10.1038/nprot.2010.166.

Keiblinger, K. M. et al. (2012) ‘Soil metaproteomics - Comparative evaluation of protein extraction protocols’, Soil Biology and Biochemistry. doi: 10.1016/j.soilbio.2012.05.014.

Lide, D. R. (2005) ‘CRC Handbook of Chemistry and Physics, Internet Version 2005’, CRC Press, Taylor and Francis Boca Raton FL. doi: 10.1016/0165-9936(91)85111-4.

Molday, R. S., Englander, S. W. and Kallen, R. G. (1972) ‘Primary Structure Effects on Peptide Hydrogen Exchange’, Biochemistry.

Sachsenberg, T. et al. (2015) ‘MetaProSIP: Automated inference of stable isotope incorporation rates in proteins for functional metaproteomics’, Journal of Proteome Research. doi: 10.1021/pr500245w.

Senko, M. W., Beu, S. C. and McLaffertycor, F. W. (1995) ‘Determination of monoisotopic masses and ion populations for large biomolecules from resolved isotopic distributions’, Journal of the American Society for Mass Spectrometry. doi: 10.1016/1044-0305(95)00017-8.

Starke, R., Kermer, R., et al. (2016) ‘Bacteria dominate the short-term assimilation of plant-derived N in soil’, Soil Biology and Biochemistry, 96. doi: 10.1016/j.soilbio.2016.01.009.

Starke, R., Keller, A., et al. (2016) ‘Pulsed C<inf>2</inf>-Acetate Protein-SIP Unveils Epsilonproteobacteria as Dominant Acetate Utilizers in a Sulfate-Reducing Microbial Community Mineralizing Benzene’, Microbial Ecology, 71(4). doi: 10.1007/s00248-016-0731-y.

Starke, R. et al. (2020) ‘Tracing incorporation of heavy water into proteins for species-specific metabolic activity in complex communities’, Journal of Proteomics. doi: 10.1016/j.jprot.2020.103791.

Starke, R., Jehmlich, N. and Bastida, F. (2018) ‘Using proteins to study how microbes contribute to soil ecosystem services: The current state and future perspectives of soil metaproteomics’, Journal of Proteomics. doi: 10.1016/j.jprot.2018.11.011.

Taubert, M. et al. (2012) ‘Protein-SIP enables time-resolved analysis of the carbon flux in a sulfate-reducing, benzene-degrading microbial consortium’, ISME Journal. doi: 10.1038/ismej.2012.68.

Taubert, M. et al. (2018) ‘Tracking active groundwater microbes with D 2 O labelling to understand their ecosystem function’, Environmental Microbiology. doi: 10.1111/1462-2920.14010.

